# The HPV8 E6 protein targets the Hippo and Wnt signaling pathways as part of its arsenal to restrain keratinocyte differentiation

**DOI:** 10.1101/2023.06.22.546184

**Authors:** Sharon C. Wu, Miranda Grace, Karl Munger

## Abstract

Infections with β-genus HPVs cause hyperplastic cutaneous lesions. In individuals with the rare hereditary skin disease, epidermodysplasia verruciformis, such lesions can progress to cutaneous squamous cell carcinomas (cSCC). β-HPV infections may also underlie cSCC development in chronically immunosuppressed individuals. Despite their prevalence and disease association, these viruses are not as well studied as the cancer-associated high-risk α-genus HPVs. HPV-associated lesions are characterized by a marked expansion of dividing, basal-like, poorly differentiated viral cells that contain viral genomes. This reflects the ability of HPVs to inhibit epithelial cell differentiation which is likely driven by the need to establish and maintain long-term viral infections in basal-like epithelial cells. Remarkably, the β-HPVs accomplish this by targeting different cellular effectors than the α-genus HPVs. It was previously reported that the HPV8 E6 protein restrains epithelial cell differentiation by inhibiting Notch and TGF-β signaling. Here we report that the HPV8 E6 protein can subvert Hippo signaling by activating TEAD transcriptional programs that inhibit the expression of keratinocyte differentiation markers. Moreover, we determined that HPV8 E6 can interfere with gene expression programs triggered by Wnt signaling by binding to the β-catenin-associated transcriptional co-activator BCL9L and that this also serves to restrain the expression of epithelial differentiation markers. Hence the HPV8 E6 protein has evolved a remarkably large array of mechanisms to subvert the differentiation program of the infected epithelial cells.

**IMPORTANCE:** Human papillomaviruses (HPVs) infect basal epithelial cells and cause a dramatic expansion of basal-like, proliferative cells. This reflects the ability of papillomaviruses to delay keratinocyte differentiation thereby maintaining aspects of the basal cell identity of persistently infected cells. This may enable papillomaviruses to establish and maintain long-term infections in squamous epithelial tissues. Previous work has revealed that the ability of β-HPV8 E6 protein to inhibit NOTCH and TGF-β signaling importantly contributes to this activity. Here we present evidence that HPV8 E6 also subverts Hippo and Wnt signaling and that these activities also aid in restraining keratinocyte differentiation.

## INTRODUCTION

Papillomaviruses are a large family of epitheliotropic, non-enveloped viruses with double-stranded circular DNA genomes that have been identified in almost all vertebrate species. The >400 fully sequenced human papillomavirus genotypes (HPVs) have been phylogenetically classified into five genera, α, β, γ, μ, and ν (1, 2). The high-risk α-HPVs infect oral and anogenital tract mucosal sites and are well-studied etiological agents of cancer. They cause most cervical cancer cases and a large proportion of other anogenital cancers as well as oropharyngeal carcinomas (3). The β-HPVs predominantly infect cutaneous epithelia. The β-HPV5 and 8 were initially identified as the causal agents of cutaneous squamous cell carcinoma (cSCC) in patients with the rare genetic disorder Epidermodysplasia Verruciformis (EV) (4, 5), and β-HPVs have also been associated with cSCCs that arise as a frequent complication in long-term immunosuppressed organ transplant patients (6). Although β-HPVs are detected in actinic keratosis, a hyperproliferative precursor lesion to cSCCs, these viruses are detected only in a small fraction of the tumor cells. Hence, the link between β-HPV infections and cSCC development, particularly in the general population, remains tenuous (7–10).

The E6 and E7 proteins are the main carcinogenic drivers of high-risk α-HPVs. The two proteins are consistently expressed in cancers and their expression is necessary for tumor maintenance. They lack intrinsic enzymatic activities, and function via binding to and usurping the activities of host cellular regulatory proteins. High-risk α-HPV E6 and E7 canonically interact with and promote the degradation of the p53 and retinoblastoma (pRB) tumor suppressors, respectively (11). While the E7 proteins of the β-HPV5 and HPV8 can bind pRB, they do so with reduced affinity compared to high-risk α-HPV16 E7 and there is no evidence for pRB destabilization (12, 13). High-risk α-HPV E6 proteins target p53 for degradation by binding to the ubiquitin ligase, E6AP (UBE3A) (14, 15). The cellular targets of the high-risk α-and EV-associated β-HPV5 and HPV8 E6 proteins diverge. HPV5 and HPV8 E6 proteins do not target p53 for degradation (16) but bind to the Notch transcriptional coactivator, MAML1, and the transforming growth factor β (TGF-β) signaling pathway transcriptional co-activators SMAD2 and SMAD3, thereby dampening Notch and TGF-β signaling (17–22).

To identify host cell protein targets of HPV8 E6, we performed affinity purification coupled with mass spectrometry (AP/MS). These experiments uncovered additional, previously unknown HPV8 E6 interactors in the Hippo and Wnt signaling pathways. Like Notch and TGF-β, Hippo and Wnt are developmental signaling pathways that are dysregulated in various cancers (23–25). The Hippo signaling pathway was discovered in *Drosophila* as a regulator of organ size control. The pathway is highly conserved in mammals and consists of a cytoplasmic kinase cascade and a nuclear transcriptional module consisting of the TEA domain (TEAD) family of transcription factors. The two modules are connected by nuclear translocation of the transcriptional co-activators, Yes-Associated Protein (YAP) and Transcriptional Coactivator with PDZ-binding motif (TAZ) (23, 26). The Hippo pathway controls cell proliferation in response to diverse microenvironmental cues and cellular stress signals. One of the first signals recognized to regulate Hippo signaling was cell crowding (27). In response to such triggers, a cytoplasmic kinase cascade is activated, and YAP or TAZ is phosphorylated, retained in the cytoplasm, and targeted for proteasomal degradation. This halts TEAD-mediated transcription in the nucleus (23, 26). It was previously reported that HPV8 E6 can inhibit Hippo signaling in response to cytokinesis failure by decreasing the activation of the LATS kinases which control the degradation of YAP (28).

The *Wnt1* gene was originally discovered as the locus of a proviral insertion and a putative oncogenic driver in mouse mammary tumor virus (MMTV)-induced tumors (29). The Wnt pathway is conserved in all metazoans and plays a central role in body axis development and polarity across phyla (25, 30, 31). The Wnt proteins are ∼40 kDa secreted proteins that upon binding to their receptors inhibit glycogen synthase kinase 3β (GSK3β) and casein kinase 1α (CK1α) mediated phosphorylation and degradation of β-catenin. This enhances β-catenin nuclear translocation where it interacts with members of the T-cell factor (TCF)/lymphoid enhancer-binding factor (LEF) transcription factor family and promotes target gene expression by scaffolding transcriptional coactivators such as PYGO, BCL9, and BCL9L, and epigenetic factors such as p300 (32). Some cutaneous HPV E6 proteins, including HPV8 E6, were shown to increase WNT signaling, albeit at markedly lower levels than high-risk α-HPV E6 proteins (33).

The mechanisms by which β-HPVs appropriate Hippo and Wnt signaling components for functions relevant to their life cycles, and whether and how this may contribute to their oncogenic activities remains sparsely studied. Here, we present evidence that HPV8 E6 can bind transcriptional components of the Hippo and Wnt signaling pathway. This results in the activation of the transcriptional output of Hippo signaling and inhibition of Wnt-signaling mediated transcriptional responses. We also show that subversion of these two pathways importantly contributes to the ability of β-HPV E6 proteins to delay epithelial differentiation.

## RESULTS

### The HPV8 E6 protein can associate with components of the Hippo and Wnt signaling pathways

To identify cellular protein interactors of HPV8 E6, AP/MS analyses were performed after transiently transfecting HCT116 colon carcinoma cells with CMV-driven expression vectors encoding amino or carboxyl terminally FLAG/HA epitope-tagged HPV8 E6 proteins. A list of “high confidence” interacting proteins was obtained after subtracting common contaminants. In addition to the previously reported regulators of the Notch and TGF-β receptor signaling pathways (18, 19) we identified several components of the Hippo and Wnt signaling pathways (Table S1; Fig 1).

**Figure 1:**
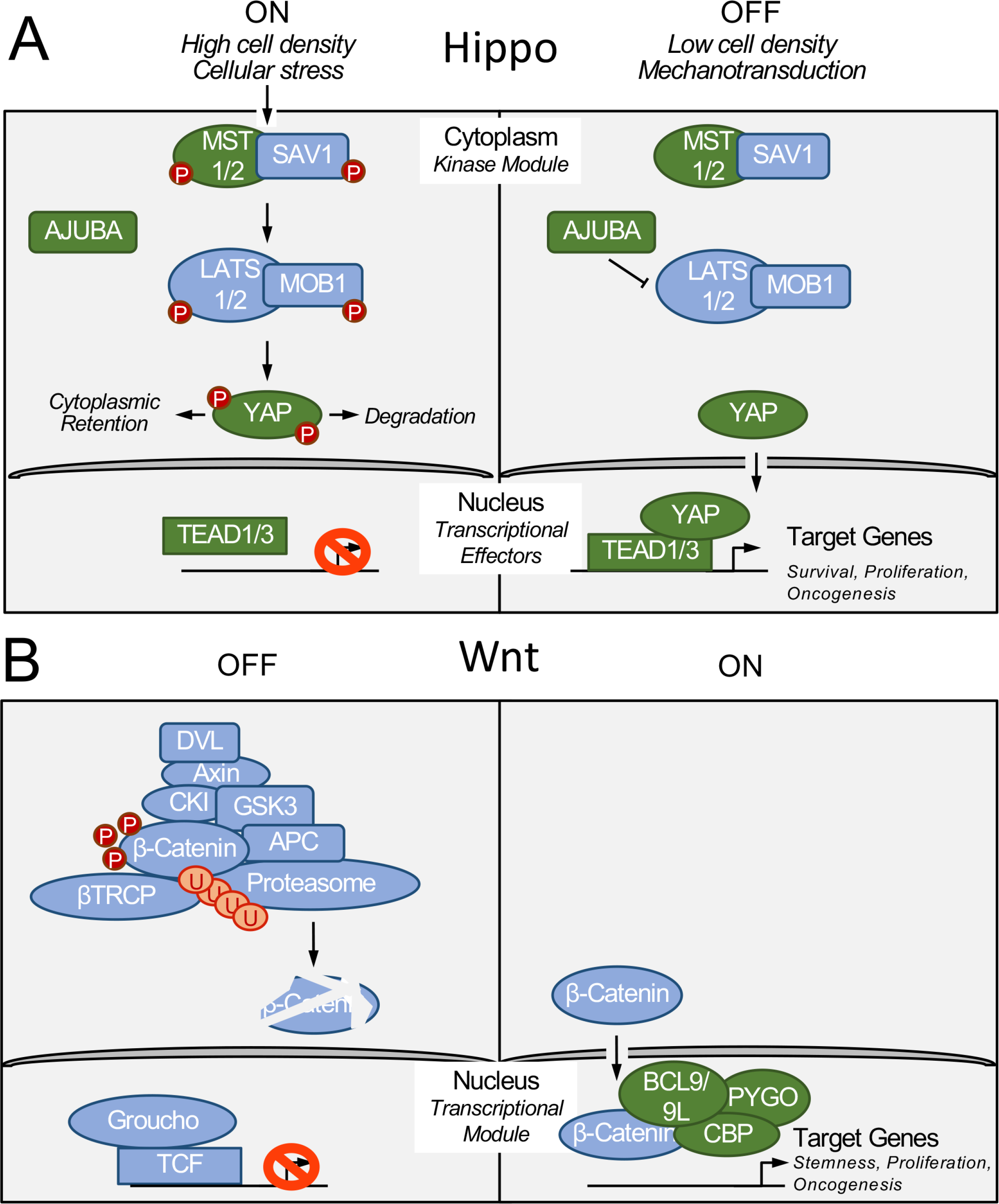
Schematic representation of the Hippo (A) and Wnt (B) signaling pathways. Signaling components identified by affinity purification/mass spectrometric analyses of HPV8 E6-associated cellular proteins are shown in green. See Table S1 for a complete list of candidate proteins and text for details.

Identified components of the Hippo signaling pathway included the cytoplasmic MST2 (STK3) kinase, and AJUBA, a negative regulator of YAP phosphorylation that is also located in the cytoplasm. Interestingly, we also identified the DNA-binding transcriptional effector of Hippo signaling, TEAD1, as a potential HPV8 E6 interacting protein. A smaller number of peptides corresponding to the related TEAD3 protein as well as the transcriptional co-activator YAP were also detected (Table S1; Fig 1A). Members of the Wnt signaling pathway that were identified by AP/MS as putative HPV8 E6 interactors included the transcriptional cofactors BCL9L, BCL9, PYGO2, and the p300-related CREB binding protein (CREBBP) (Table S1; Fig. 1B). The HCT116 cells that were used for these experiments are hemizygous for p300 and express the protein at low levels (34) and unlike in previous experiments with other cell types we did not detect p300 in our AP/MS experiments with HPV8 E6 (20, 35).

### HPV8 E6 interacts with transcriptional effectors of Hippo signaling

Given that the effects of HPV8 E6 on the cytoplasmic kinase cascade have been investigated previously (28), we focused our studies on investigating whether and how the β-HPV8 E6 protein might affect the nuclear effectors of Hippo signaling.TEAD1 was corroborated as an HPV8 E6 interaction partner via co-immunoprecipitation using either HCT116 colon cancer epithelial cells transiently transfected with FLAG/HA-tagged HPV8 E6 (Fig. 2A) or telomerase immortalized normal oral keratinocytes (NOKs) with stable expression of FLAG/HA-tagged HPV8 E6 (Fig. 2B). Co-precipitation with MAML1 was used as a positive control. It is noted that under the experimental conditions that were used for these experiments, TEAD1 co-precipitated less efficiently with HPV8 E6 than MAML1. We identified an HPV8 E6 mutant, HPV8 E6 K_136_N that is defective for interacting with TEAD1. This mutant can still efficiently interact with MAML1 and SMAD3 (Fig. 2C), suggesting that the TEAD1 binding site on HPV8 E6 is distinct from the sequences required for MAML1 or SMAD3 binding. Unfortunately, however, this mutant does not accumulate to the same level as the wild-type HPV8 E6 protein and hence we did not use it in any biological experiments. A weak interaction with the Hippo pathway transcriptional coactivator Yes-associated protein 1 (YAP1), was only detected in HPV8 E6 transiently transfected HCT116 cells but not in NOKs with stable HPV8 E6 expression. Thus, HPV8 E6 efficiently interacts with TEAD1 but not with YAP (Fig 2D).

**Figure 2:**
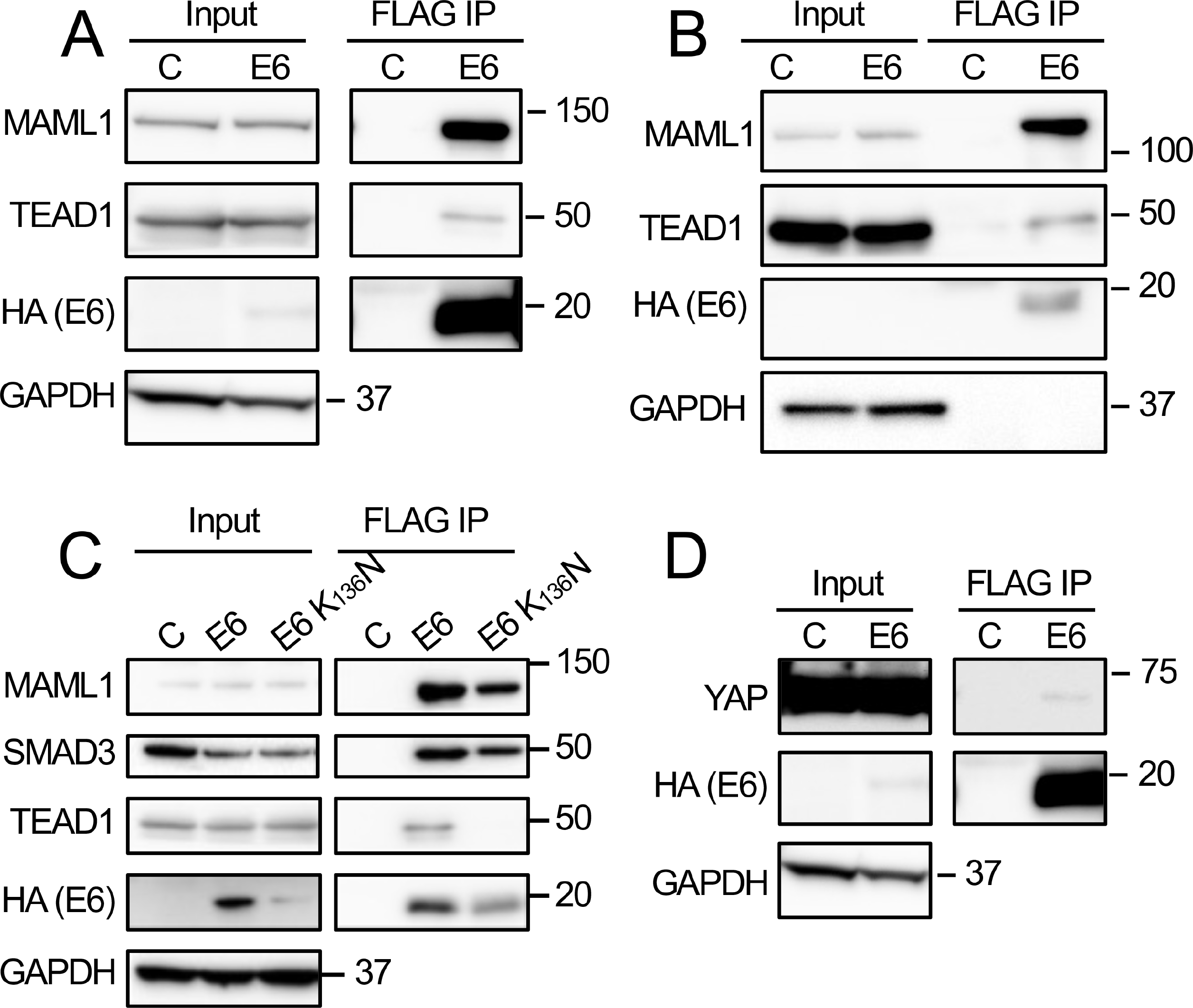
The HPV8 E7 protein can associate with nuclear effectors of Hippo signaling. FLAG immunoprecipitations were performed with extracts from HCT116 colon carcinoma cells transiently transfected with N-terminally FLAG/HA-tagged HPV8 E6 (E6), a TEAD binding defective N-terminally FLAG/HA-tagged HPV8 E6 K136N mutant, or a control vector (C) **(A, C, D)** or with extract from telomerase immortalized normal human oral keratinocytes (NOKs) with stable expression of an N-terminally FLAG/HA-tagged HPV8 E6 protein (E6) or control vector transduced NOKs (C) **(B)**. MAML1, TEAD1, SMAD3, YAP, and HPV8 E6 levels were assessed by western blotting. GAPDH was used as a loading control. MAML1 co-IP was used as a positive control.

### HPV8 E6 activates YAP/TEAD-mediated transcriptional programs

To determine whether HPV8 E6 may affect TEAD-mediated transcriptional responses, we first used a luciferase reporter assay. An effector plasmid where the DNA-binding domain of the yeast transcription factor Gal4 is fused to TEAD1 (Gal4dbd-TEAD1), was co-transfected with a reporter plasmid expressing firefly luciferase under the control of the upstream activation sequence of Gal4 (Gal4UAS-Luc) and an HPV8 E6 expression plasmid or a YAP expression plasmid as a positive control. A renilla luciferase expression vector was included to control for transfection efficiency. Transfection of HPV8 E6 led to a consistent statistically significant, ∼3-fold, increase in reporter activity as compared to an ∼10-fold increase upon YAP co-transfection. Co-transfection of HPV8 E6 in addition to YAP did not cause an additional significant increase in reporter activity (Fig 3A). Next, we assayed the expression of the YAP/TEAD target genes amphiregulin (*AREG*) and angiomotin-like 2 (*AMOTL2*) (36, 37) in NOKs with stable expression of an HA-FLAG-tagged HPV8 E6 expression vector or a control plasmid. These two genes were chosen from a study that identified core Hippo target genes in a variety of carcinomas using data from The Cancer Genome Atlas (TCGA) (36). YAP/TEAD target gene expression is influenced by cell density and is lower at high cell density (27). HPV8 E6 significantly enhanced the expression of *AREG* at high but not at cell low density (Fig 3B). HPV8 E6 also enhanced the expression of *AMOTL2* mRNA at high density, although this increase did not reach statistical significance (Fig 3C). Hence, HPV8 E6 can activate TEAD-mediated transcriptional responses.

**Figure 3:**
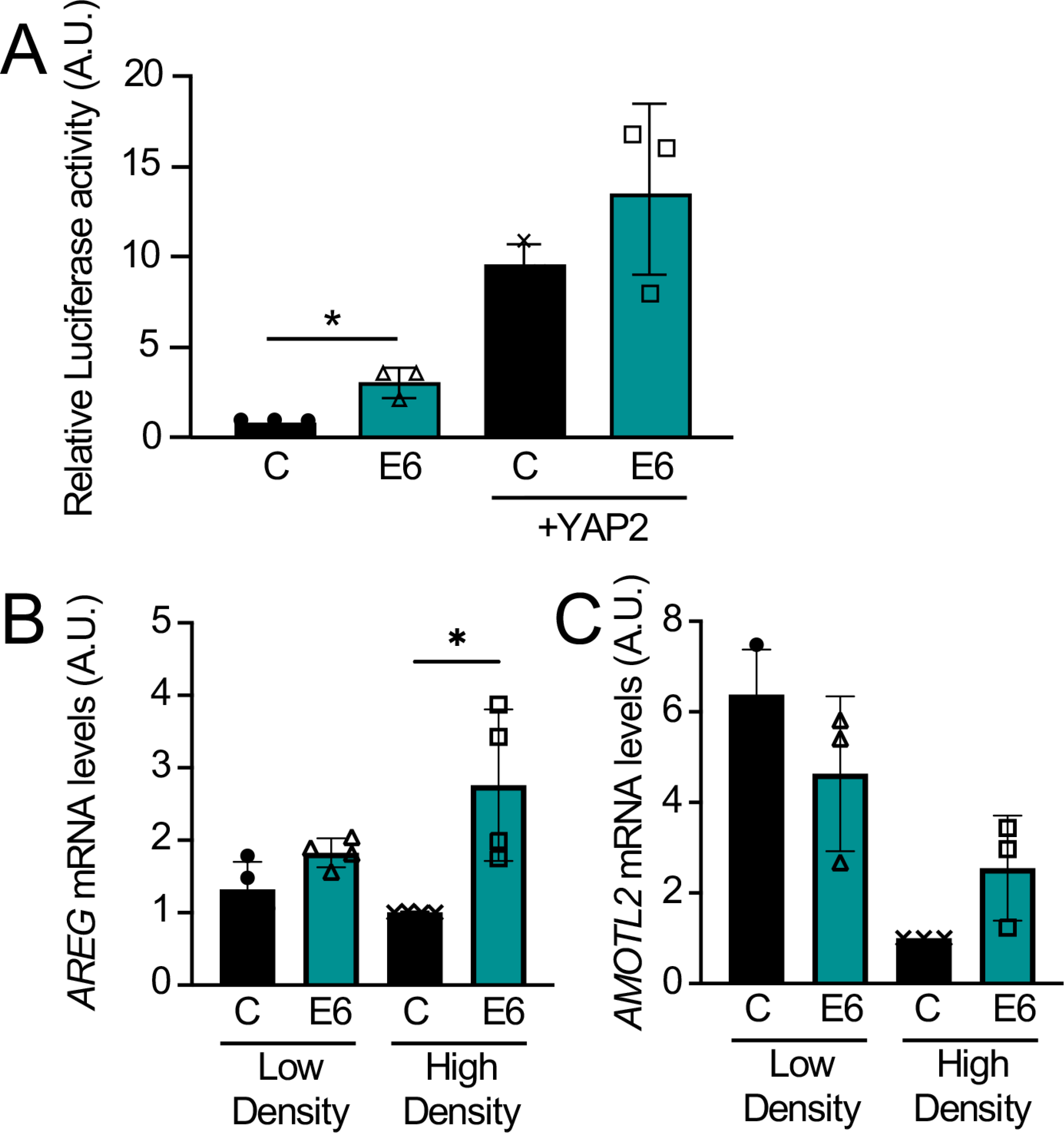
HPV8 E6 activates YAP/TEAD-mediated transcription: HCT116 cells were transfected with 100 ng Gal4dbd-TEAD1 and Gal4uas-Firefly luciferase vectors, 4 ng Renilla luciferase internal control, 10 ng of FLAG-YAP2 or empty vector control, and 200 ng of N-terminally FLAG/HA-tagged HPV8 E6 (E6) or empty vector as a control (C). Firefly luciferase activity normalized to Renilla luciferase was quantified. Data represent averages and standard deviations from three independent experiments. Statistical significance was assessed by the Kruskal-Wallis test. * denotes p ≤ 0.05 **(A)** Telomerase immortalized normal human oral keratinocytes (NOKs) with stable expression of an N-terminally FLAG/HA-tagged HPV8 E6 protein (E6) or control vector transduced NOKs (C) were grown to ∼50% confluency (Low Density) or 100% confluency (High Density) before harvest. Expression of *AREG* and *AMOTL2* mRNAs was assessed via qRT-PCR. Data were normalized to RPLP0 as a housekeeping gene. Data shown are means from 4 (AREG) or 3 (AMOTL2) independent experiments. A 2-way ANOVA with Sidak’s multiple comparisons test was performed to assess statistical significance. ** denotes p ≤ 0.01. **(B and C).**

### Loss of TEAD1 or TEAD3 attenuates the ability of HPV8 E6 to inhibit keratinocyte differentiation

After providing evidence that E6 can enhance TEAD-mediated transcription, we wanted to investigate the biological relevance of the interaction of HPV8 E6 with TEAD family members. One well-known biological activity of HPV8 E6 is to delay keratinocyte differentiation. To determine whether the interaction of HPV8 E6 with TEAD family members may contribute to the inhibition of keratinocyte differentiation, TEAD1 or TEAD3 were depleted by RNAi in HPV8 E6-expressing or control NOKs before subjecting the cells to calcium-mediated differentiation. Transfection with a non-targeting control siRNA was used as a control. The mRNA levels of keratin K10 (*KRT10*) and filaggrin (*FLG*), markers for early and late stages of keratinocyte differentiation, respectively, were determined by quantitative reverse transcription PCR (Q RT-PCR). As expected, HPV8 E6 expressing NOKs showed marked defects in differentiation as evidenced by lower *KRT10* and *FLG* mRNA levels compared to control vector transduced NOKs. Consistent with an earlier publication (38), depletion of TEAD1 or TEAD3 did not significantly affect the differentiation-induced expression of *KRT10* or *FLG* mRNA expression in control vector transduced NOKs (Figs 4A, B). In contrast, however, depletion of *TEAD3*, but not *TEAD1*, significantly rescued *KRT10* expression in HPV8 E6-expressing NOKs (Fig. 4A). Moreover, depletion of *TEAD1* but not *TEAD3*, significantly rescued *FLG* expression in E6-expressing NOKs (Fig. 4B). In contrast, individual depletion of TEAD1 or TEAD3 only had minimal effects on involucrin *(IVL)* expression in control of HPV8 E6 expressing NOKs (Fig 4C). Q RT-PCR analyses confirmed that the TEAD1-specific siRNA pool did not affect TEAD3 mRNA levels and vice versa (Fig. S1A). These results suggest that HPV8 E6 can act through TEAD1 and TEAD3 to inhibit the expression of specific keratinocyte differentiation-associated genes. We also performed analogous experiments in hTERT-immortalized human foreskin keratinocytes (iHFKs). In these cells, individual depletion of TEAD1 or TEAD3 alone did not rescue differentiation marker expression in HPV8 E6-expressing iHFKs (Fig S2). It is thus possible that in keratinocytes derived from specific anatomic locations, the TEAD isoforms exhibit functional redundancy, and the loss of a single TEAD family member is insufficient to rescue the expression of differentiation markers in HPV8 E6 expressing cells. Additional experiments will be necessary to parse this potential context-dependent redundancy.

**Figure 4:**
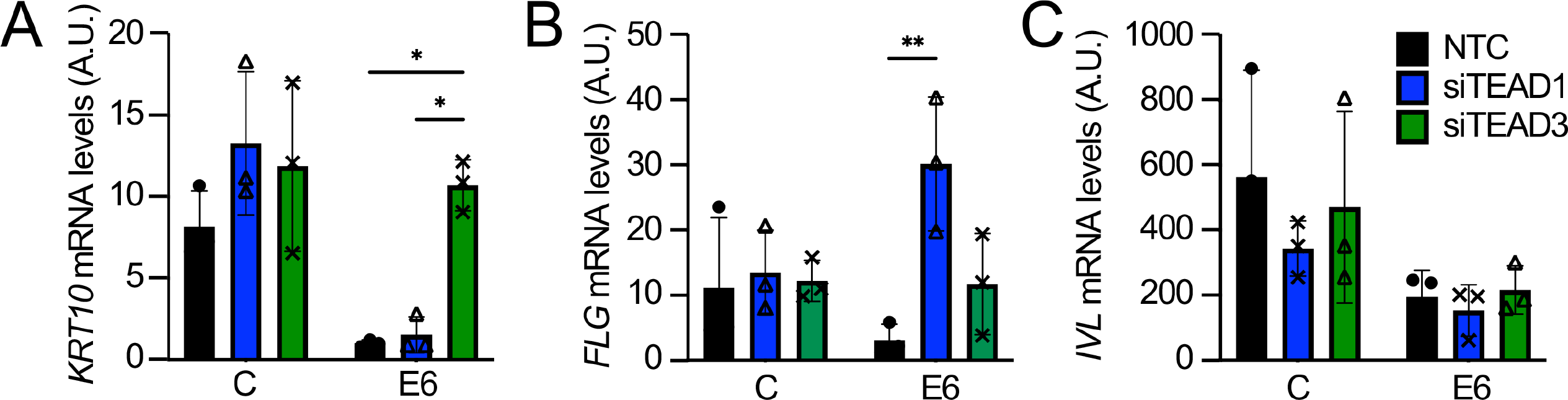
TEAD1 or TEAD3 depletion attenuates the ability of HPV8 E6 to inhibit keratinocyte differentiation. Telomerase immortalized normal human oral keratinocytes (NOKs) with stable expression of an N-terminally FLAG/HA-tagged HPV8 E6 protein (E6) or control vector transduced NOKs (C) were grown to 100% confluency and switched to 10% FBS-containing Dulbecco’s modified Eagle Medium (DMEM) to induce differentiation and transfected with TEAD1 (siTEAD1), TEAD3 (siTEAD3) onTARGETplus SMARTpools, or a non-targeting control siRNA pool (NTC), and grown for 4 days. Keratin K10 (*KRT10*) **(A)**, Filaggrin (*FLG*) **(B)**, and Involucrin (*IVL*) **(C)** mRNA levels were assessed by qRT-PCR. Gene expression for each condition at 4 days was normalized to expression in Day 0 control NOKs before the medium switch. Data were normalized to RPLP0 as the housekeeping gene. The data shown are means from 3 independent experiments. P-values were calculated using 2-way ANOVA with Sidak’s multiple comparisons test. ** = p ≤ 0.01

### HPV8 E6 interacts with the Wnt transcriptional co-activator BCL9L

The components of the Wnt pathway, BCL9, BCL9L, PYGO, and CREBBP that we identified by AP/MS as candidate HPV8 E6 interactors are all part of the Wnt nuclear effector complex. The transcriptional coactivator and histone acetyltransferase CREBBP acts as transcriptional co-activator in various signaling pathways (39). Both high-risk α and β HPV E6 proteins have been reported to interact with CREBBP and/or the highly related p300 protein (20, 40, 41), and β HPV8 E6 has been reported to cause p300 degradation (42). In the Wnt signaling pathway, p300 has been reported to bind β-catenin and promote β-catenin-mediated transcription and oncogenic transformation (43). In other studies, however, CREBBP and p300 have also been reported to repress Wnt transcriptional output (44, 45). We focused our experiments on Wnt signaling on the nuclear Wnt co-activator BCL9L because, to our knowledge, it has never been reported as an HPV E6 interactor and, unlike BCL9 and PYGO2, it was identified as an interactor with both N- and C-terminally tagged HPV8 E6 and at a high peptide count (Table S1). The interaction between HPV8 E6/BCL9L interaction was confirmed via co-immunoprecipitation in HCT116 cells transiently transfected with FLAG/HA-tagged HPV8 E6 (Fig 5A) as well as in telomerase immortalized human foreskin keratinocytes (iHFKs) with stable expression of FLAG/HA-tagged HPV8 E6 (Fig 5B).

**Figure 5:**
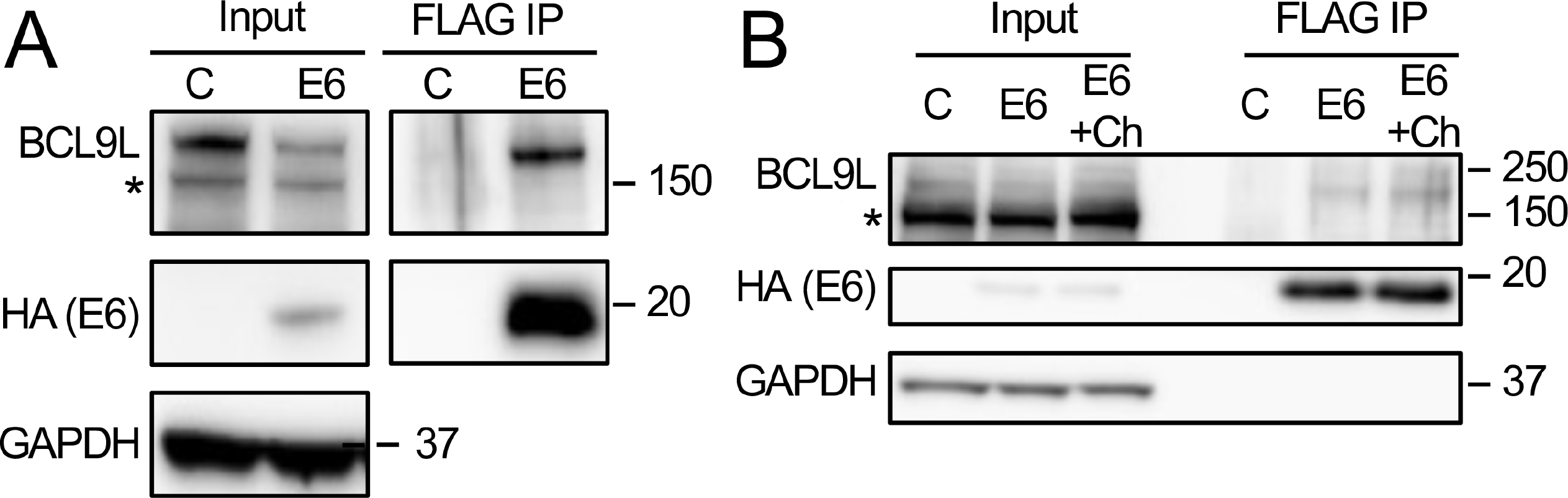
HPV8 E6 interacts with the Wnt transcriptional co-activator BCL9L. FLAG immunoprecipitations were performed with extracts from HCT116 colon carcinoma cells transiently transfected with N-terminally FLAG/HA-tagged HPV8 E6 (E6) or a control vector (C) **(A)** or with extracts from telomerase immortalized normal human foreskin keratinocytes (iHFKs) with stable expression of an N-terminally FLAG/HA-tagged HPV8 E6 protein (E6) that were left untreated or treated with the Wnt activator CHIR99021 (+Ch) or control vector transduced iHFKs (C) **(B)**. BCL9L and HPV8 E6 levels were assessed via western blotting. GAPDH was used as a loading control.

To determine whether the interaction of HPV8 E6 with BCL9L was affected by Wnt activation, we treated the control and HPV8 E6 expressing cells with the widely used pharmacological inducer of Wnt signaling, the glycogen synthase kinase 3 (GSK3) inhibitor CHIR99021 (46). GSK3 restrains canonical Wnt signaling by phosphorylating β-catenin, thereby targeting it for proteasomal degradation, and GSK3 inhibition results in the stabilization and enhanced nuclear translocation of β-catenin (47). These experiments showed that activation of Wnt signaling does not affect the interaction of HPV8 E6 with BCL9L (Fig 5B).

### HPV8 E6 inhibits the transcriptional output of Wnt signaling

To determine whether HPV8 E6 may modulate the transcriptional output of Wnt signaling, we performed luciferase reporter assays in HCT116 cells with the Wnt-responsive Super 8x TOPFlash reporter plasmid, where *Firefly* luciferase expression is regulated by eight TCF/LEF binding sites. The Super 8x FOPFlash reporter, which has 8 mutant TCF/LEF binding sites was used as a control. Co-transfection of HPV8 E6 significantly decreased Super 8x TOPFlash reporter activity compared to control (Fig 6A). We next examined the expression of the Wnt target gene *Axin2* (47) in HPV8 E6 expressing and control vector transduced iHFKs. Baseline *Axin2* expression was low in both cell lines. A 6-hour treatment with 3 µM CHIR99021 induced *Axin2* expression in control iHFKs and this response was significantly lower in the HPV8 E6 expressing iHFKs (Fig 6B). Hence, HPV8 E6 can inhibit the transcriptional output of canonical Wnt signaling.

**Figure 6:**
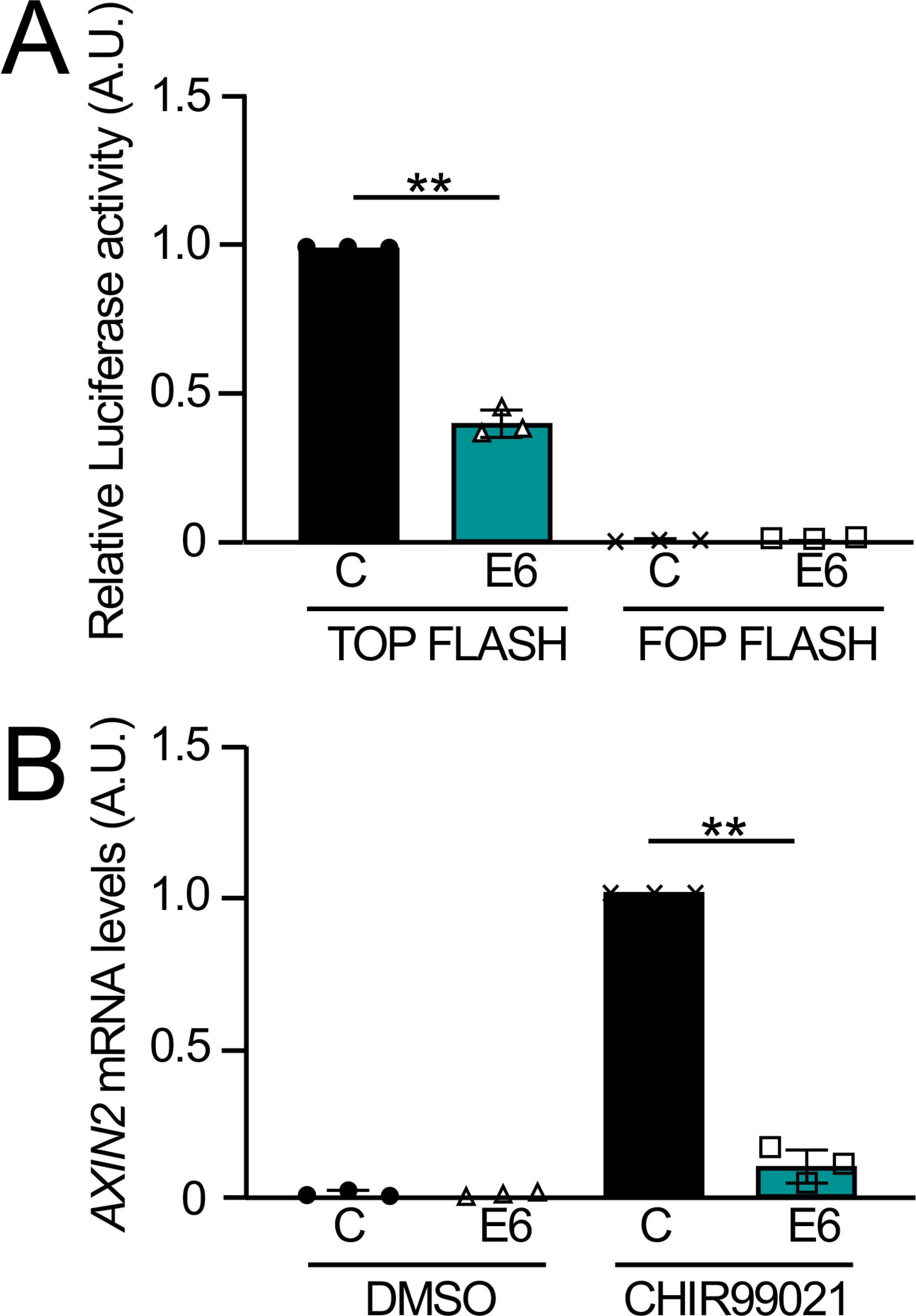
HPV8 E6 inhibits Wnt transcriptional activation. HCT116 cells were transfected with 200 ng TOP-FLASH or FOP-FLASH reporter vectors; 4 ng Renilla luciferase internal control vector; and 50 ng of FLAG/HA-tagged HPV8 E6 or empty vector control. Firefly normalized to Renilla luciferase activity was quantified. Data from 3 independent experiments are shown. An unpaired 2-tailed t-test with Welch’s correction was performed to calculate the p-value. ** = p ≤ 0.01 **(A).** Telomerase immortalized normal human foreskin keratinocytes (iHFKs) with stable expression of an N-terminally FLAG/HA-tagged HPV8 E6 protein (E6) or control vector transduced cells (C) were treated with 3 µM to the Wnt activator CHIR99021 or vehicle control (DMSO) for 6 hours. mRNA levels of the canonical Wnt pathway target gene *AXIN2* were assessed via qRT-PCR. Expression was normalized to GAPDH as the housekeeping gene. Bar graphs represent means and standard deviations from 3 independent experiments. An unpaired 2-tailed t-test with Welch’s correction was performed to calculate the p-value. ** denotes p ≤ 0.01 **(B)**.

### HPV8 E6 expression inhibits Wnt-mediated expression of keratinocyte differentiation markers

Wnt signaling has been extensively studied in the context of keratinocyte biology. On one hand, increased Wnt signaling has been linked to epithelial stem cell maintenance whereas in other experiments Wnt signaling was shown to contribute to epithelial differentiation (48). Hence, we investigated whether pharmacological Wnt activation may modulate the differentiation of human keratinocytes. iHFKs were induced to differentiate by switching near confluent cultures from low-calcium keratinocyte serum-free medium (KSFM) to Dulbecco’s Modified Eagle Medium (DMEM) with 10% fetal bovine serum for four days. To study the effect of Wnt signaling, the cells were concurrently treated with 3 µM CHIR99021 or DMSO as a control. As expected, our differentiation protocol resulted in a significant increase in mRNA expression of the early differentiation marker keratin K10 (*KRT10*) in the control vector transduced but not in the HPV8 E6 expressing cells. Concurrent treatment with the pharmacological Wnt activator, CHIR99021, caused a further significant increase in *KRT10* mRNA expression (Fig 7A). Similarly, involucrin (*IVL*) mRNA levels were more prominently induced in control cells than in HPV8 E6 expressing cells, and expression further significantly increased in response to pharmacological Wnt activation (Fig 7B). While the mRNA expression of the late differentiation marker filaggrin (*FLG*) was not markedly induced by our differentiation protocol, CHIR99021 treatment caused a marked increase in *FLG* mRNA expression in control but not in the HPV8 E6 expressing cells (Fig 7C). Hence, pharmacological Wnt activation stimulates the expression of differentiation markers in normal human foreskin keratinocytes cells and HPV8 E6 can interfere with this process.

**Figure 7:**
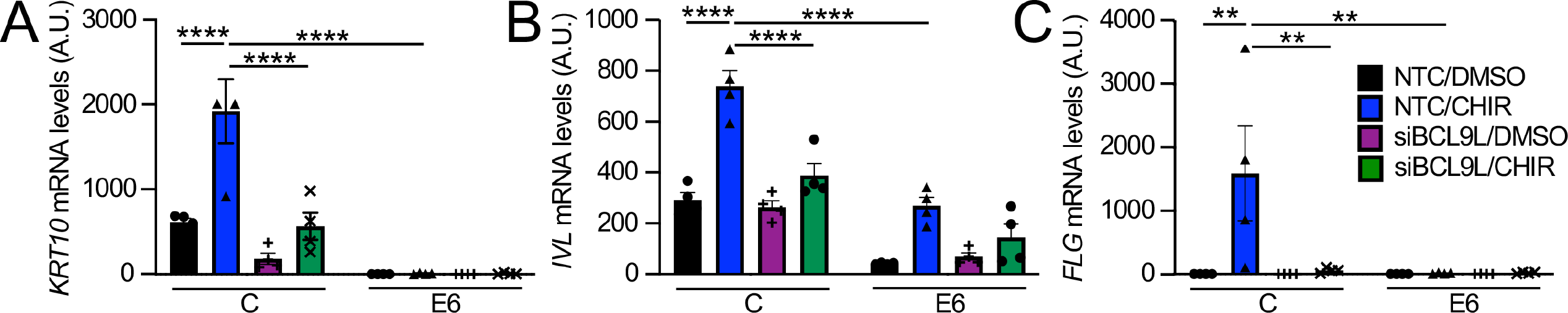
CHIR99021 promotes calcium differentiation in a BCL9L-dependent manner, which is blocked by HPV8 E6. Telomerase immortalized human foreskin keratinocytes (iHFKs) with stable expression of an N-terminally FLAG/HA-tagged HPV8 E6 protein (E6) or control vector transduced NOKs (C) were grown to 100% confluency and switched to 10% FBS-containing Dulbecco’s modified Eagle Medium (DMEM) to induce differentiation, transfected with a BCL9L onTARGETplus SMARTpools or non-targeting control siRNA pool (NTC), concurrently treated with 3 µM CHIR99021 or vehicle (DMSO), and grown for 4 days. Keratin K10 (*KRT10*) **(A)**, Involucrin (*IVL*) **(B)**, and Filaggrin (*FLG*) **(C)** mRNA levels were assessed by qRT-PCR. Gene expression for each condition at 4 days was normalized to expression in Day 0 control iHFKs before the medium switch. GAPDH was used as the housekeeping gene. Bar graphs represent averages and standard deviations from four independent experiments. P-values were calculated using 2-way ANOVA with Sidak’s multiple comparisons tests. *= p ≤ 0.05 ** = p ≤ 0.01 *** = p ≤ 0.001 **** = p ≤ 0.0001

### Wnt signaling-induced expression of keratinocyte differentiation markers is dependent on BCL9L

Given the ability of HPV8 E6 to bind the Wnt-coactivator, BCL9L, (Fig 5A, B), we next investigated whether the observed ability of Wnt signaling to induce expression of keratinocyte differentiation markers (Fig. 7) was dependent on BCL9L. To experimentally address this question, HPV8 E6 expressing and control vector transduced iHFKs were transfected with a BCL9L targeting siRNA pool (siBCL9L) or a non-targeting control (NTC) before they were subjected to calcium-mediated differentiation either in the presence or absence of CHIR99021, as described above. These experiments revealed that BCL9L depletion significantly reduced induction of *KRT10*, *IVL*, and *FLG* mRNA expression in response to activation of Wnt signaling (Fig 7). These experiments show that pharmacological activation of Wnt signaling promotes the expression of keratinocyte differentiation markers through a BCL9L-dependent mechanism and that BCL9L depletion interferes with HPV8 E6-mediated inhibition of keratinocyte differentiation.

These experiments were also performed in NOKs. The expression of differentiation markers is somewhat different in NOKs than in iHFKs as evidenced by the induction of filaggrin in NOKs (Fig S3C) but not in iHFKs (Fig 7C) by our differentiation protocol. Hence it is not surprising that the effects of CHIR99021 on NOK differentiation were more varied. Similar to the iHFKs, CHIR99021 treatment enhanced *K10*, and more prominently, *FLG mRNA* expression in control NOKs, and this effect was attenuated in HPV8 E6-expressing NOKs. Surprisingly, CHIR99021 treatment decreased *IVL* mRNA expression in control and E6 expressing NOKs. Loss of BCL9L attenuates *K10* expression in CHIR99021-treated control as well as E6-expressing NOKs, whereas the effect on *FLG* mRNA expression was more dramatic in the E6-expressing NOKs. In contrast, BCL9L depletion did not affect *IVL* mRNA expression (Fig S3). Nonetheless, these results are consistent with our model that pharmacological Wnt pathway activation can enhance the expression of some differentiation markers under high calcium conditions, and HPV8 E6 can blunt this effect. BCL9L may contribute to the Wnt pathway activation-mediated expression of some keratinocyte differentiation markers but the transcriptional effectors necessary for differentiation may be different in keratinocytes derived from different anatomic locations.

## DISCUSSION

The expansion of poorly differentiated, basal-like cells is a histopathological hallmark of papillomavirus-associated lesions. This reflects the ability of papillomaviruses to delay keratinocyte differentiation presumably to maintain the basal cell identity of infected cells, which enables papillomaviruses to establish long-term persistent infections in squamous epithelia. Papillomaviruses have evolved multiple mechanisms to delay keratinocyte differentiation. The β-HPV8 and MmuPV1 E6 proteins inhibit NOTCH and TGF-β signaling, by binding the MAML1 co-activator and the SMAD2 and SMAD3 proteins, respectively (49). Here we have identified two additional pathways, Hippo and Wnt signaling, that are targeted by the β-HPV8 E6 protein to inhibit keratinocyte differentiation.

Aberrant Hippo signaling in epithelial cells as a consequence of hyperactive YAP is a potent oncogenic driver and triggers rapid onset and progression of oral and cervical squamous cell carcinoma in mouse models (32, 50). A previous study showed that HPV8 E6 can affect the cytoplasmic kinase module to aberrantly activate Hippo signaling-dependent gene transcription in response to cytokinesis failure (28). Our study reveals that HPV8 E6 also targets TEAD transcription factors, which function as nuclear effectors of Hippo signaling. Previous studies have shown that simultaneous depletion of TEAD1 and TEAD3 in primary human keratinocytes attenuated cell proliferation and caused decreased expression of the differentiation markers, filaggrin, and loricrin. Individual depletion of TEAD1 or TEAD3, however, did not affect proliferation, terminal differentiation, or expression of classical Hippo target genes (38). Consistent with these results, we found that TEAD1 or TEAD3 depletion did not affect the expression of early and late differentiation genes in normal human keratinocytes. Intriguingly, however, TEAD1 and TEAD3 depletion differentially affected the expression of keratinocyte differentiation markers in HPV8 E6 expressing keratinocytes where differentiation is inhibited. Depletion of TEAD3, but not TEAD1 caused increased expression of keratin K10, a marker of the spinous layer. In contrast, TEAD1 depletion caused increased expression of the granular layer marker filaggrin much more potently than TEAD3 depletion. Hence TEAD1 and TEAD3 may have distinct targets during the earlier phases of differentiation but may work in concert to promote the differentiation program during later stages of differentiation. Thus, HPV8 E6 may co-opt different TEAD family members to dysregulate early and late differentiation genes. Alternatively Taken together, these results suggest that HPV8 E6 can uncouple the cytoplasmic kinase cascade from its nuclear transcriptional effectors to aberrantly sustain the transcriptional output of the Hippo signaling pathway even under homeostatically inhibitory conditions.

The mechanism by which HPV8 E6 may usurp the TEADs to sustain the transcriptional output of Hippo signaling to delay differentiation is unknown. The fact that HPV8 E6 only very weakly interacts with YAP by co-immunoprecipitation and does not specifically increase the nuclear population of YAP may suggest that HPV8 E6 can affect TEAD-dependent transcription independent of YAP. Two models are possible: HPV8 E6 may promote the expression of TEAD target genes that encode proteins that can suppress the expression of differentiation markers, or HPV8 E6 may recruit repressive machinery to TEAD to inhibit the transcriptional activation of keratinocyte differentiation markers. Identification of and functional experiments with stably expressed TEAD-binding defective HPV8 E6 mutants that maintain binding to other known HPV8 E6 interactors will further illuminate whether and how interaction with TEAD family members contributes to the inhibition of differentiation.

The high-risk mucosal α-HPV E7 proteins subvert Hippo signaling by targeting the non-receptor tyrosine phosphatase and tumor suppressor, PTPN14, for degradation through the associated UBR4 ubiquitin ligase (51, 52). E7/UBR4-mediated PTPN14 degradation cause activation of YAP which stimulates YAP/TEAD-mediated gene transcription (53). This has been linked to the inhibition of keratinocyte differentiation and maintenance of basal cell identity (53, 54). The β-HPV E7 proteins can also bind UBR4 and PTPN14, although both UBR4 and PTPN14 are bound less efficiently by β-HPV E7 than by the high-risk α-HPV E7, and β-HPV E7 proteins have minimal effects on the steady-state level of PTPN14 (20, 35, 52, 55). Hence it remains to be determined if HPV8 E7 also contributes to dysregulating TEAD-mediated transcriptional programs.

A recently published study with EBV has shown that the EBV LMP1 protein subverts Hippo signaling through multiple mechanisms which lead to phosphorylation and nuclear translocation of YAP and the related TAZ protein. This is required for EBV-mediated induction of proliferation, inhibition of differentiation, and induction of the epithelial-to-mesenchymal transition (EMT) in NOK cells (56).

Our finding that HPV8 E6 can inhibit transcriptional programs downstream of Wnt activation was surprising since previous studies have reported evidence that Wnt/β-catenin signaling is upregulated in lesions and cancers caused by mucosal high-risk α-HPVs (57). There are fewer studies that investigate the effects of β-HPVs on Wnt signaling, but one study suggests β-HPV E6 proteins may also activate Wnt signaling albeit less efficiently than HPV16 E6 (33). Our results, however, indicate that HPV8 E6 can inhibit the transcriptional output of canonical Wnt signaling both by using a TCF/LEF-responsive luciferase reporter and by assessing mRNA expression of the canonical target gene, AXIN2 (47). The β-catenin-associated transcriptional coactivator BCL9L was confirmed as a previously unknown interactor of HPV8 E6 and we showed that HPV8 E6-mediated inhibition of Wnt signaling is dependent on BCL9L. Pharmacological activation of canonical Wnt signaling with the GSK3β inhibitor CHIR99021, which causes stabilization and enhanced nuclear translocation of β-catenin triggers expression of keratinocyte differentiation markers, and this is dependent on BCL9L.

The mechanism by which BCL9L may promote keratinocyte differentiation in response to calcium, however, is not known. Despite the model that Wnt activation drives cancer progression by promoting stemness and oncogenic processes and thus, the Wnt-cancer connection appears to be more nuanced, requiring a “just right”, intermediate level of activation for oncogenesis (58). Subversion of Hippo and Wnt signaling by HPV8 E6 may not only cause inhibition of keratinocyte differentiation but also affects other cellular activities. Hippo signaling is activated in tetraploid cells that result from cytokinesis failure through activation of the LATS2 kinase which results in p53 stabilization and YAP and TAZ inactivation (59). It was shown that HPV8 E6 can stabilize p53 which leads to tolerance of genomic instability (60). Later work showed that LATS phosphorylation and consequently Hippo signaling was inhibited in HPV8 E6 expressing cells which promoted the tolerance of failed cytokinesis and centrosome replication errors thereby increasing the frequency of multinucleated cells which can lead to the development of aneuploidy (28, 61). The HPV8 E6 and E7 proteins are potent mutagens. They can interfere with DNA double-strand break repair (62) and generate chromosomal rearrangements and micronuclei formation (63). The ability of the β-HPVs to subvert genomic integrity and allow for the maintenance of such cells in the proliferative pool is particularly interesting since these viruses are readily detected in precursor lesions but not in frank cancers (9, 10). Hence β-HPVs may contribute to skin carcinogenesis through a “hit-and-run” mechanism and subversion of genomic integrity and inhibition of mechanisms that normally eliminate such cells from the proliferative pool is a plausible explanation of how even the transient expression of viral sequences of cells could mechanistically contribute to carcinogenesis (64). Given that UV exposure promotes chromosomal instability in keratinocytes (65), β-HPVs likely evolved mechanisms to promote host cell tolerance to genotoxic insults to ensure its maintenance and propagation in a UV-exposed niche. Interestingly, loss of BCL9L was also shown to promote aneuploidy tolerance (66). One may speculate that the interaction of HPV8 E6 with BCL9L may also contribute to the increased survival of cells with chromosomal anomalies. Subversion of Hippo signaling by HPV8 E6 is also predicted to cause alterations in cell adhesion (67) and to enhance survival under conditions of limiting glucose concentrations (68). Because pharmacological Wnt activation can upregulate interferon β (IFN-β) expression thereby enhancing innate immune signaling (69), HPV8 E6 mediated inhibition of Wnt signaling may also serve to down-modulate innate immune responses.

There is evidence for crosstalk between Hippo, Wnt, and Notch signaling. In the mouse epidermis, Notch1 has been found to repress Wnt signaling, and Notch1^-/-^ mice exhibit increased β-catenin levels (70). Notch1 is expressed in the suprabasal layers of the mouse epidermis whereas β-catenin expression is restricted to the basal epidermal layers, suggesting that Notch1 might repress β-catenin in differentiating keratinocytes (70). The inhibition of Notch signaling by HPV8 E6 is predicted to increase β-catenin levels, and hence HPV8 E6 might suppress Wnt transcriptional programs to counter this. Moreover, activation of TEAD-mediated transcription in epidermal cells inhibits Notch signaling and differentiation, thereby preserving stem cell-like traits and basal cell identity (71). Whether activation of TEAD-mediated transcription by HPV8 E6 contributes to the observed repression of Notch-mediated differentiation or whether Notch signaling drives the expression of keratinocyte differentiation markers that we observed upon depletion of TEAD family members in HPV8 E6-expressing cells remains to be determined.

The β HPV genus is heterogeneous and comprises 5 species. This study focuses on the HPV8 E6 protein, a β-1 species member. However, host protein interactions and thus pathway perturbations by HPV8 E6 may not be generalizable to β HPVs within other species (41, 72). Hence it will be interesting to determine whether E6 proteins encoded by other β-HPVs, or more generally other HPVs, may similarly interact with TEADs and/or BCL9L to subvert the Hippo and Wnt signaling pathways.

## MATERIALS AND METHODS

### Cell Culture

Human foreskin keratinocytes immortalized with human telomerase (hTERT HFK Cl 398 - iHFK) were a gift from Aloysius Klingelhutz (73). iHFKs and telomerase-immortalized normal oral keratinocytes (NOKs) (74) were cultured in keratinocyte serum-free media (KSFM) supplemented with bovine pituitary extract and human epidermal growth factor (Thermo Fisher). HCT116 human colon carcinoma lines were obtained from ATCC and maintained following their recommendations. Transient transfection of HCT116 cells was performed as previously described (75). HPV8 E6 expressing keratinocytes were generated via lentiviral transduction as reported previously (18). CHIR99021 (Cayman) was used at 3 μM. Calcium-mediated differentiation was induced by switching keratinocytes from KSFM to 10% fetal bovine serum-containing Dulbecco’s modified Eagle medium (DMEM) at 100 % confluency.

### Luciferase Assays

M50 Super 8x TOPFlash (Addgene plasmid #12456) and M51 Super 8x FOPFlash (Addgene plasmid #12457) were gifts from Randall Moon (76). Gal4DBD-TEAD1, Gal4UAS-Luc, and CMV-YAP plasmids were gifts from Dr. Brian Schaffhausen. pRLgk Renilla was a gift from Dr. Elliott Kieff. Cells were transfected using FuGene6 (Promega). For the Hippo experiments, the following plasmids were transfected: 100 ng Gal4dbd-TEAD1; 100 ng Gal4uas-Luc; 4 ng pRLgk Renilla; 200 ng CMV-N-HPV8 E6 or CMV-N empty vector; and 10 ng CMV-YAP or empty vector. For the Wnt experiments in HCT116 cells, the following plasmid amounts were transfected: 200 ng M50 Super 8x TOPFlash or M51 Super 8x FOPFlash; 4 ng pRLgk Renilla; and 50 ng of CMV-N-HPV8 E6 or CMV-N empty vector. Lysates were harvested 2 days hours following transfection and reporter assays were performed using the Dual-Luciferase Reporter Assay (Promega).

### RNA isolation and quantitative reverse transcription PCR

RNA isolation, reverse transcription, quantitative PCR, and analysis were described previously (77). The following primers were used:

*RPLP0:* 5’-TGGTCATCCAGCAGGTGTTCGA-3’ (Fwd) and 5’-ACAGACACTGGCAACATTGCGG-3’ (Rev) (sequences from Origene);

*AREG:* 5’-GCACCTGGAAGCAGTAACATGC-3’ (Fwd) and 5’-GGCAGCTATGGCTGCTAATGCA-3’ (Rev) (sequences from Origene);

*AMOTL2:* 5’-AGTGAGCGACAAACAGCAGACG-3’ (Fwd) and 5’-ATCTCTGCTCCCGTGTTTGGCA-3’ (Rev) (sequences from Origene);

*GAPDH:* 5’-GATTCCACCCATGGCAAATTC-3’ (Fwd) and 5’-TGGGATTTCCATTGATGACAAG-3’ (Rev)

*Involucrin:* 5’-TGCCTGAGCAAGAATGTGAG-3’ (Fwd) and 5’-TGCTCTGGGTTTTCTGCTTT-3’ (Rev);

*Filaggrin:* 5’-AAAGAGCTGAAGGAACTTCTG-3’ (Fwd) and 5’-AACCATATCTGGGTCATCTGG-3’ (Rev)

*K10:* 5’-GCAAATTGAGAGCCTGACTG-3’ (Fwd) and 5’-CAGTGGACACATTTCGAAGG-3’ (Rev);

*TEAD1:* 5’-CCTGGCTATCTATCCACCATGTG-3’ (Fwd) and 5’-TTCTGGTCCTCGTCTTGCCTGT-3’ (Rev) (sequences from Origene);

*TEAD3:* 5’-AGGCAGTAGATGTGCGCCAGAT-3’ (Fwd) and 5’-TCCTGGATGGTGCTGTTGAGGT-3’ (Rev) (sequences from Origene);

*BCL9L:* 5’-CCGCTCTACCACAATGCCATCA-3’ (Fwd) and 5’-CTGAGTTCAGGTGCATCTGGCT-3’ (Rev) (sequences from Origene);

*BCL9:* 5’-TCCAGCTCGTTCTCCCAACTTG-3’ (Fwd) and 5’-GATTGGAGTGAGAAAGTGGCTGG-3’ (Rev) (sequences from Origene)

*PYGO1:* 5’-GGTTAGGAGGACCAGGTGTACA-3’ (Fwd) and 5’-AGCAGCCACTAGATGGTCAGAG-3’ (Rev) (sequences from Origene)

### Immunoprecipitation, REAP nuclear/cytoplasmic fractionation, Western Blot

EBC buffer (50 mM Tris HCl pH 8.0, 150 mM NaCl, 0.5% NP40, 0.5 mM EDTA) was used for cell lysis. Immunoprecipitations were conducted using Anti-FLAG M2 Affinity Gel (Sigma). The immunoprecipitated proteins and the whole cell input were run on NuPAGE 4-12% BisTris gels. Proteins were transferred onto a Polyvinylidene fluoride (PVDF) membrane (Millipore) and blocked with 5% nonfat dry milk in TBST (20 mM Tris-HCl, 137 mM NaCl, 0.1% Tween 20, pH 7.6) or TNET (20 mM Tris-HCl, 100 mM NaCl, 5 mM EDTA, 0.1% Tween 20, pH 7.5) buffer (77). Blots were probed with the following primary antibodies: TEAD1 (D9X2L, 1:1000, CST12292S; GAPDH (1:400-500, MAB374); HA (1:200, sc805 - SW33, SW38); HA (1:5000, ab9110); YAP (1:1000, CST4912S); BCL9L (1:200, AF4967) SMAD3 (1:1000, C67H9 CST #9523); hnRNP (1:1000, ab10294); Tubulin (5 µg/mL, ab18251). Blots were washed with TBST or TNET and probed with horseradish peroxidase-conjugated secondary antibodies. This was followed by further washing, application of chemiluminescent substrate, and imaging using the G:Box Chemi-XX6 imager with Genesys software. The TEAD-binding defective HPV8 E6 K_136_N mutant was made using the QuikChange II site-directed mutagenesis kit (Agilent). Nuclear/cytoplasmic fractionation was performed using the previously published Rapid, Efficient, and Practical cellular fractionation protocol (78).

### siRNA transfection

Keratinocytes were transfected with onTARGETplus SMARTpool, consisting of a mixture of four different siRNAs, against human TEAD1, TEAD3, or BCL9L Non-targeting siRNA SMARTpools were used as a control. Transfection was performed using RNAIMax transfection reagent (Sigma).

## ACKNOWLEDGEMENTS

We thank Dr. Al Klingelhutz (University of Iowa) for providing telomerase immortalized human foreskin keratinocytes, and Drs. Marta Gaglia, Phil Hinds, Alexander Poltorak, and members of the Munger and Lambert Groups for stimulating discussions and suggestions throughout this work. Special thanks to Dr. Elizabeth White for her valuable insights and comments on this manuscript. Supported by PHS grants R01 CA228543 (K.M.) and T32 GM008448 (SCW). K.M. dedicates this paper to the memory of Dr. Massimo Tommasino.

---

**Figure S1: Assessment of TEAD1, TEAD3, and BCL9L depletion in the experiments shown in Figure 4 (A) and Figure 7 (B).**

**Figure S2: TEAD1 or TEAD3 depletion attenuates the ability of HPV8 E6 to inhibit keratinocyte differentiation in telomerase-immortalized human foreskin keratinocytes (iHFKs).** Experiments were performed as described in Figure 4. Data shown are averages of 2 (*KRT10*, *FLG*) and 3 (*IVL*) independent experiments each performed in triplicate.

**Figure S3: TEAD1 or TEAD3 depletion attenuates the ability of HPV8 E6 to inhibit keratinocyte differentiation in telomerase-immortalized normal human oral keratinocytes (NOKs).** Experiments were performed as described in Figure 7. Data shown are averages of 3 (*IVL*) and 4 (*KRT10*, *FLG*) independent experiments each performed in triplicate.

**Table S1: Interactors identified with AP/MS**

List of HPV8 E6 interactors identified via affinity purification/mass spectrometry in HCT116 colon cancer cells transfected with HPV8 E6 tagged with FLAG and HA at the C terminus (“HPV8 CE6”) or N-terminus (“HPV8 NE6”). Both number of unique peptides (“Unique”) and total peptides (“Total”) are shown.

